# The efficacy of CB-103, a first-in-class transcriptional Notch inhibitor, in preclinical models of breast cancer

**DOI:** 10.1101/2023.07.06.547830

**Authors:** Michele Vigolo, Charlotte Urech, Sebastien Lamy, Giulia Monticone, Jovanny Zabaleta, Fokhrul Hossain, Dorota Wyczechowska, Luis Del Valle, Ruth O’Regan, Lucio Miele, Rajwinder Lehal, Samarpan Majumder

## Abstract

**Background:** Notch signaling has been shown to mediate treatment resistance and support cancer stem cells (CSCs) in endocrine-resistant estrogen receptor positive (ER+) and triple negative breast cancers (TNBCs). The clinical development of GSIs, first generation Notch inhibitors, has been hindered by lack of Notch specificity and dose-limiting toxicity. Here we describe the safety and efficacy of a first-in-class, clinical stage, orally available small molecule pan-Notch inhibitor, CB-103. Due to its unique mode of action, CB-103 doesn’t induce GI toxicities noted with GSIs. There is a critical need for effective, safe, targeted therapies for patients with endocrine-refractory metastatic breast cancer. Recently approved targeted therapies for TNBC are only effective for a subset of patients. Moreover, GSI-resistant, constitutively activating Notch1 or Notch2 mutations are observed in ∼10% of TNBC. Our study elucidating the synergy of CB-103 with fulvestrant and paclitaxel in preclinical models of both hormone-refractory ER+ BC and TNBC respectively provides a novel and unique opportunity to address major unmet therapeutic needs.

**Methods:** CB-103 was screened in combination with a panel of anti-neoplastic drugs. We evaluated the anti-tumor activity of CB-103 with fulvestrant in ESR1-mutant (Y537S), endocrine-resistant BC xenografts. In the same model, we examined anti-CSC activity in mammosphere formation assays for CB-103 alone or in combination with fulvestrant or palbociclib. We also evaluated the effect of CB-103 plus paclitaxel on primary tumors and CSC in GSI-resistant TNBC model HCC1187. Comparisons between groups were performed with two-sided unpaired Student’s *t*-test. One-way or two-way ANOVA followed by Tukey’s post analysis was performed to analyze in vivo efficacy study results.

**Results:** CB-103 showed synergism with fulvestrant in ER+ cells and with paclitaxel in TNBC cells. CB-103 combined with fulvestrant or paclitaxel potently inhibited mammosphere formation in both models. Combination of CB-103 and fulvestrant significantly reduced tumor volume in an ESR1-mutant, endocrine-resistant BC model. In a GSI-resistant TNBC model, CB-103 plus paclitaxel significantly delayed tumor growth compared to paclitaxel alone.

**Conclusions:** Our data indicate that CB-103 is an attractive candidate for clinical investigation in endocrine resistant, recurrent breast cancers with biomarker-confirmed Notch activity in combination with SERDs and/or CDKis and in TNBCs with biomarker-confirmed Notch activity in combination with taxane-containing chemotherapy regimens.

## Background

Despite recent advances in the treatment of metastatic hormone receptor (HR)-positive, i.e. ER+ and/or PR+ breast cancers, endocrine resistance ultimately develops in all cases. Additionally, a significant proportion of HR-positive breast cancers exhibit intrinsic endocrine resistance. In the metastatic setting, available second line therapies are moderately effective and can have significant toxicities, and third-line therapies are generally largely ineffective. Thus, there is a critical need for new therapies that can circumvent endocrine resistance to further improve the outcomes of HR-positive disease. TNBC is an aggressive and heterogeneous breast cancer subtype that accounts for 15−25% of all breast cancer diagnoses in Western countries (1). Patients with early TNBC have a two– to three-fold higher risk of disease recurrence and death in the first three years after diagnosis than patients with non-TNBC (2). TNBC disproportionately affects young premenopausal women and African-American (AA) women. Only recently the US FDA approved a few targeted therapies for TNBC patients (3). However, the clinical benefits of these agents are restricted to limited subgroups of patients, and chemotherapy remains the mainstay of treatment. Notch signaling is involved in chemoresistance in breast cancer (4) and specifically in paclitaxel resistance (5).

The role of Notch signaling in endocrine-resistance is well-established (6–8). Meta-analyses of tumor molecular landscapes and a number of pathology studies show that Notch expression and/or activity are associated with risk of recurrence in HR-positive breast cancers (9). Estrogen deprivation or tamoxifen cause activation of Notch1 in ER-positive breast cancer cells, and Notch inhibition dramatically increased the efficacy of tamoxifen in MCF-7 xenografts, causing tumor regression (10). We showed that Notch1 can activate ERα-dependent transcription in the absence of estrogen, evading estrogen deprivation (11) and that Protein Kinase C (PKC)α, a known marker of endocrine resistance, induces endocrine resistance in HR-positive breast cancer cells via Notch4 (12). Mutations in the ESR1 gene affecting the hormone binding domain of its product ERα are associated with Notch activation in breast cancer cell lines (13).

Notch signaling activation is also associated with TNBC tumor growth, CSC maintenance and expansion, tumor invasiveness, and metastasis (6, 14, 15). Importantly, the appearance of Notch-dependent cancer stem-like cells (CSC) was shown to be responsible for resistance to mTOR inhibitors in TNBC (16). Therefore, the Notch signaling pathway has been the object of intense pre-clinical and clinical investigation as a possible therapeutic target in breast cancer (6). Inhibition of Notch in tumors where the pathway is active can potentially produce growth inhibition, apoptosis and angiogenesis, while simultaneously inhibiting CSC (6). The first generation of Notch inhibitors tested in the clinic were GSIs (6). However, the development of this class of drugs has been hindered by low specificity (γ-secretase has nearly 150 known substrates) (17) and dose-dependent, on-target gastrointestinal toxicity (6, 18). Two GSIs remain in late clinical development for desmoid tumors and salivary adenoid cystic carcinoma (ACC) (6). However, despite some encouraging results in early phase studies in endocrine-resistant breast cancer (19), the clinical development of GSIs in breast cancer has languished, underscoring the need for less toxic and more specific next-generation Notch inhibitors. Importantly, at least in TNBC a significant fraction of cases harbor constitutively activating Notch mutations that do not require γ-secretase cleavage (3), underscoring the need to inhibit Notch through targets other than γ-secretase.

CB-103 is a first-in-class, non-GSI, orally available, highly specific protein-protein interaction (PPI) inhibitor that interferes with the interaction between the active intracellular domains of Notch receptors (NICD) and the CSL transcription factor complex (6, 20). CB-103 blocks both ligand-dependent and ligand-independent Notch transcription without effecting the myriad other γ-secretase substrates. Its unique mode of action suggests that CB-103 would be effective against the GSI-resistant, truncated forms of Notch1 or Notch2 generated by genetic rearrangements associated with ∼10% of TNBC (3), in addition to other Notch-dependent breast tumors. The safety and efficacy of CB-103 in Notch-dependent advanced or metastatic solid or hematological malignancies have been investigated in a multi-center phase I/II clinical trial (Clinicaltrials.gov: <u>NCT03422679</u>) (3). In earlier clinical trials, CB-103 has been safe and well-tolerated, showing minimal gastrointestinal toxicity and preliminary efficacy signals in solid tumors and leukemias (6, 18). We first tested CB-103 in a PDX model of ER+BC with wild type estrogen receptor α gene (ESR1). In this model, CB-103 showed activity similar to fulvestrant but not synergy with fulvestrant. However, when the same combination was tested in an ESR1 mutated xenograft model, CB-103 showed synergy with fulvestrant. Here we present evidence that CB-103 in combination with fulvestrant arrested the growth of mouse xenografts from a patient-derived endocrine-resistant model carrying an ESR1 Y537S mutation. CB-103 alone and in combinations with either fulvestrant or palbociclib showed potent anti-mammosphere activity in the same model. Furthermore, we document potent tumor growth inhibition and delayed tumor relapse when CB-103 was combined with paclitaxel in the GSI-resistant HCC1187 TNBC cell line model. In the same HCC1187 model, CB-103 alone and in combination with paclitaxel showed potent anti-mammosphere activity, while paclitaxel alone had none. Our results, along with early phase clinical trial results, provide strong rationale for testing CB-103 in Notch-dependent endocrine-resistant breast and/or TNBC.

## Methods

### CB-103 combination screening

Human cancer cell lines HCC1187 and MCF-7 were obtained from the laboratory of Freddy Radtke and ATCC, respectively. Cells were cultured under mycoplasma-free conditions at 37°C in RPMI-1640 media supplemented with 10% FCS (HCC1187 cells) or DMEM media supplemented with 10% FCS (MCF-7 cells). Cells were seeded at a density of 1500 cells/well in 384-well plate format and cultured in 40 ml of growth media. Cells were treated with a combination of CB-103 (concentration range from 150 nM to 10 μM) and chemical compounds listed in Table 1 (concentration range from 75 nM to 20 μM) for 72 hours. The concentration range used for each compound was determined based on individual IC_50_ values, *i.e.* concentration range selected flanks IC_50_ value of each compound. Each of the compounds were combined with CB-103 at different concentrations generating an 8X10 matrix in duplicates. Compounds were prepared as stock solution at 10 mM in pure DMSO. Purity was checked by LC-MS and was above 90% for all solutions. To create 384-well working plates, different volumes of stock solutions were plated into 384 well plates by using ECHO 550 acoustic dispenser (Labcyte) to generate the final concentration of interest for each drug. To determine growth kinetics, alamarBlue® (Invitrogen) was added to each well and incubated for 4 hours. alamarBlue® readout was taken using Infinite® F500 (Tecan) plate reader. All volumes (cells and alamarBlue®) were dispensed using Multidrop Combi dispenser using a standard cassette (Speed MEDIUM). Synergistic interaction analysis was performed using SynergyFinder software (http://bioconductor.org/packages/release/bioc/html/synergyfinder.html or https://synergyfinder.fimm.fi/) as described by Ianevsky et al. (21).

**Table 1.** – Most synergistic area score of drug combinations in TNBC and ER+BC cells. >0: synergistic effect; 0, no combination effect; <0, antagonist effect.

|  | Most synergistic area score |  |
| --- | --- | --- |
| Drug combination | ER+ BC (MCF-7) | TNB (HCC1187) |
| PD 0332991 (Palbociclib) + CB-103 | 15.85 | -0.01 |
| Paclitaxel + CB-103 | 24.87 | 8.73 |
| Fulvestrant + CB-103 | 24.64 | -2.49 |

### Cell lines and mammospheres

All cell lines used were authenticated by STR (short tandem repeat) analysis by an independent contract laboratory (Genetica DNA Laboratories – LabCorp). All cell lines were treated with anti-mycoplasma from time to time to ensure there was no mycoplasma contamination. The human breast cancer cell line (MCF-7) was obtained from the ATCC. The cell line was maintained in Dulbecco’s Modified Eagle Medium (DMEM; GIBCO) supplemented with 10% HiFBS, 1% Glutamax and 1% Penicillin-Streptomycin, at 37°C in 5% CO2. Other endocrine resistant cell lines used in our study were obtained from different laboratories. T47D: PKCα cells were generated in the laboratory of Dr. Debra Tonetti (UIC), by transfection with the pSPKCα plasmid by electroporation (22). Y537S was a kind gift from Dr. Matt Burow’s lab in Tulane University. Y537S mutant cell line was originally derived from WHIM20 primary tumor (23). WHIM stands for Washington University Human in Mouse PDX lines. Li et al. (23) showed estradiol-independent growth of WHIM20 which expressed an ESR1-Y537S point mutation. Secondary mammosphere cultures were established from cell lines as previously described (19) and treated with CB-103, fulvestrant, palbociclib and paclitaxel alone or in different combinations thereof at concentrations determined through pilot experiments. Mammospheres were counted after 7 days as previously described by means-Powell et al. (19).

### *In vivo* efficacy study to evaluate CB-103 vs fulvestrant and combination in therapy resistant ER+ BC

Tumor engraftment: 2×10^6^ Y537S ER^+^ ESR1 mutant BC cells were injected orthotopically beneath the 4th mammary fat pad of 40 Nude/Ovariectomized female mice, about 4 months old. No estrogen pellet was used to facilitate tumor growth. Tumor engraftment was measured using a digital caliper weekly for the duration of the experiment. Tumor volumes were calculated using the following formula: 0.5 × (Length × Width^2^). After approximately a month, we identified and selected 32 mice with tumor growth and randomized them into 4 groups of 8: Vehicle control, CB-103 alone, fulvestrant alone and combination of CB-103 and fulvestrant. Once tumors reached ∼100 mm^3^, mice were dosed subcutaneously with fulvestrant (250 mg/kg Q7D) or CB-103 (60 mg/kg QDx5) for 4 weeks. Toy et al, (24) used 200 mg/kg twice a week s.c. dosing of fulvestrant for mutant Y537S. In our hands, once a week 250 mg/kg was sufficient. The control arm received vehicle treatment (5%DMSO, 95%Castor oil). The experiment was terminated on day 29 when the vehicle treated control group reached a humane endpoint agreement with the approved animal protocol (IACUC # 3599 as per LSUHSC). Tumors were isolated from 5 best responsive mice from each group and snap frozen tumor tissues were analyzed by RNA sequencing and immunohistochemistry.

### ER+ BC PDX combination with fulvestrant

Athymic Nude-Foxn1nu (immune-compromised) female mice at 6-8 weeks of age were implanted subcutaneously in the left flank side with fragment of the PDX tumor model CTG-2308 (Champion Oncology). Tumor growth was monitored twice a week using digital calipers and the tumor volume (TV) was calculated using the formula (0.52 x [length x width^2^]). When the TV reached approximately 100-200 mm³, animals were matched by tumor size and assigned into the following dosing cohort: Vehicle control (N=10), CB-103 40mg/kg QDx4 (N=9), fulvestrant 5mg/kg Q7D (N=10) or CB-103+fulvestrant (N=9). Efficacy study terminated on day 51 of treatment when one mouse in control group reached endpoint.

### TNBC xenograft combination study with paclitaxel

NSG (immune-compromised) female mice at 6-8 weeks of age were implanted s.c. in the left flank side with 1×10^6^ HCC1187 TNBC cells. Tumor growth was monitored twice a week using digital calipers and the tumor volume (TV) was calculated using the formula (0.5 x [length x width^2^]). When the TV reached approximately 30-60 mm³, animals were matched by tumor size and dosed with either Vehicle, CB-103 60mg/kg QDx5, Paclitaxel 10mg/kg Q7D or the combination of both. In two independent experiments each dosing group switched treatment when control group reached around 1000 mm^3^. In the first case dosing was turned off for all groups. In the second case the switch occurred as following: A-Vehicle ➔ combo (N=10), B-CB-103 ➔ CB-103 (N=10), C-Paclitaxel ➔ no dosing (N=9), D-Combo ➔ no dosing (N=9), Combo ➔ CB-103 (N=10). Efficacy studies terminated respectively on day 39 and 63 after first dose, when half of group C reached endpoint in agreement with the approved animal protocol from Swiss DGAV (National license n. 33520, Cantonal license n. VD3672).

### Marginal zone B cell assay

We used 12 non-tumor bearing nude mice randomized into 3 groups-vehicle control (5%DMSO, 95%Castor oil), CB-103 40mg/kg and 60mg/kg. CB-103 and vehicle control were injected s.c. daily for 7 days. On the 8^th^ day mice were sacrificed and B cells from spleen from each of those mice were isolated and flow cytometry was performed using a cocktail of CD21, CD23 and B220 B cell specific antibody as per Lehal et al. (25).

### RNA Sequencing

RNA sequencing and analysis were performed in the Translational Genomics Core (TGC) at the Stanley S. Scott Cancer Center, LSUHSC, New Orleans, LA. RNA was isolated from CB-103, fulvestrant, combination and vehicle treated excised tumors using the Universal RNA/DNA Isolation kit (Qiagen, Germantown, MD) following the manufacturer’s protocol. Isolated RNA was quantified using a Qubit (ThermoFisher, Waltham, MA) and checked for RNA integrity on the Agilent Bio Analyzer 2100 (Agilent, Santa Clara, CA). All the analysis was done in Partek Flow and included removal of contaminants (mDNA, rDNA, tDNA) with Bowtie v2.2.5, alignment to hg19 version of the human genome using STAR v2.5.3a, and quantification of aligned reads using RefSeq Transcripts 93 (released 2020-02-03). A filter was applied to exclude transcripts with less than 5 reads in 80% of the samples (per comparison). Normalization was done with TMM and transformed by log2(+0.0001/TMM/log2). Finally, differential expression analysis was done with DESEQ2. Normalized counts were used for pathways analysis with KEGG (embedded in Partek Flow). All analyses were corrected for multiple comparisons at a false discovery rate (FDR) of 0.05. In addition, only those genes with fold change > 2.0 were considered for pathways analyses as described (26).

### Histology and Immunohistochemistry

Sections from mouse tumors were fixed in 10% buffered formalin for 24 hours. After paraffin embedding, tumors were sectioned at 4 microns in thickness and Hematoxylin and Eosin staining was performed for routing histopathological analysis. Immunohistochemistry was performed using the avidin-biotin-peroxidase methodology, according to the manufacturer’s instructions (Vectastain ABC Elite Kit, Vector Laboratories, Burlingame, CA, USA). Our modified protocol includes deparaffination in xylenes, rehydration through descending grades of ethanol up to water, non-enzymatic antigen retrieval with 0.01 M sodium citrate buffer pH 6.0 at 95 °C for 25 min, endogenous peroxidase quenching with 3% H_2_O_2_ in methanol, blocking with normal horse serum (for mouse monoclonal antibodies) or normal goat serum (for rabbit polyclonal or recombinant rabbit monoclonal antibodies) and incubation with primary antibodies overnight at room temperature in a humidified chamber. Antibodies included a mouse monoclonal anti-Ki67 (DAKO, Clone IVAK-2), 1:100 dilution), a rabbit monoclonal anti-CD31 (abcam, clone EPR17259, 1: 2000 dilution), and a rabbit polyclonal anti-Cleaved Caspase 3 (Cell Signaling, Asp175, 1:500 dilution). After rinsing in PBS, sections were incubated with anti-mouse or anti-rabbit biotinylated secondary antibodies for 1 hour, followed by incubation with avidin-biotin-peroxidase complexes for 1 hour, both at room temperature in a humidified chamber. Finally, the peroxidase was developed with diaminobenzidine (Boehringer, Mannheim, Germany) for 3 minutes, and the sections were counterstained with Hematoxylin and mounted with Permount (Fisher Scientific). Photomicrographs were taken with an Olympus DP72 Digital Camera using an Olympus BX70 microscope (Olympus, Center Valley, PA, USA).

### Statistical analysis

GraphPad Prism 9.5 software was used for the analysis of the data and graphic representations. Comparisons between groups were performed with two-sided unpaired Student’s *t*-test. One-way ANOVA analysis followed by Tukey’s test post analysis was used for mouse experiments to compare one variable among multiple groups. Two-way ANOVA analysis followed by Dunnett’s test post analysis was used for mouse experiments to compare one variable over time among multiple groups. *p* values□≤□0.05 were considered significant.

## Results

### CB-103 shows synergy with several anti-neoplastic drugs

To examine potential pharmacological interactions between CB-103 and other FDA-approved or investigational drugs, we performed in vitro assays with eight different concentrations of CB-103 in various drug combinations in both ER^+^ MCF-7 cells and the TNBC cell line HCC1187, which is Notch2-mutated and GSI-resistant. Table 1 shows the most synergistic area scores for these two models. We generated dose-response matrices (Figure 1), which showed that combinations of CB-103 and the selective estrogen receptor disruptor (SERD), fulvestrant produced robust synergy (most synergistic area score) across all the concentration ranges tested in MCF-7 (Figure 1B) but not in TNBC HCC1187 cells (Figure 1D). CB-103 also showed significant synergy with paclitaxel in both models (Figures 1A, C). Of note, CB-103 also showed robust synergy with CDK4/6 inhibitor PD 0332991 (Palbociclib) in MCF-7 but not in HCC1187 cells (Table 1). These findings support the further study of these drug combinations in ER+ breast cancer and TNBC models.

**Figure 1.**
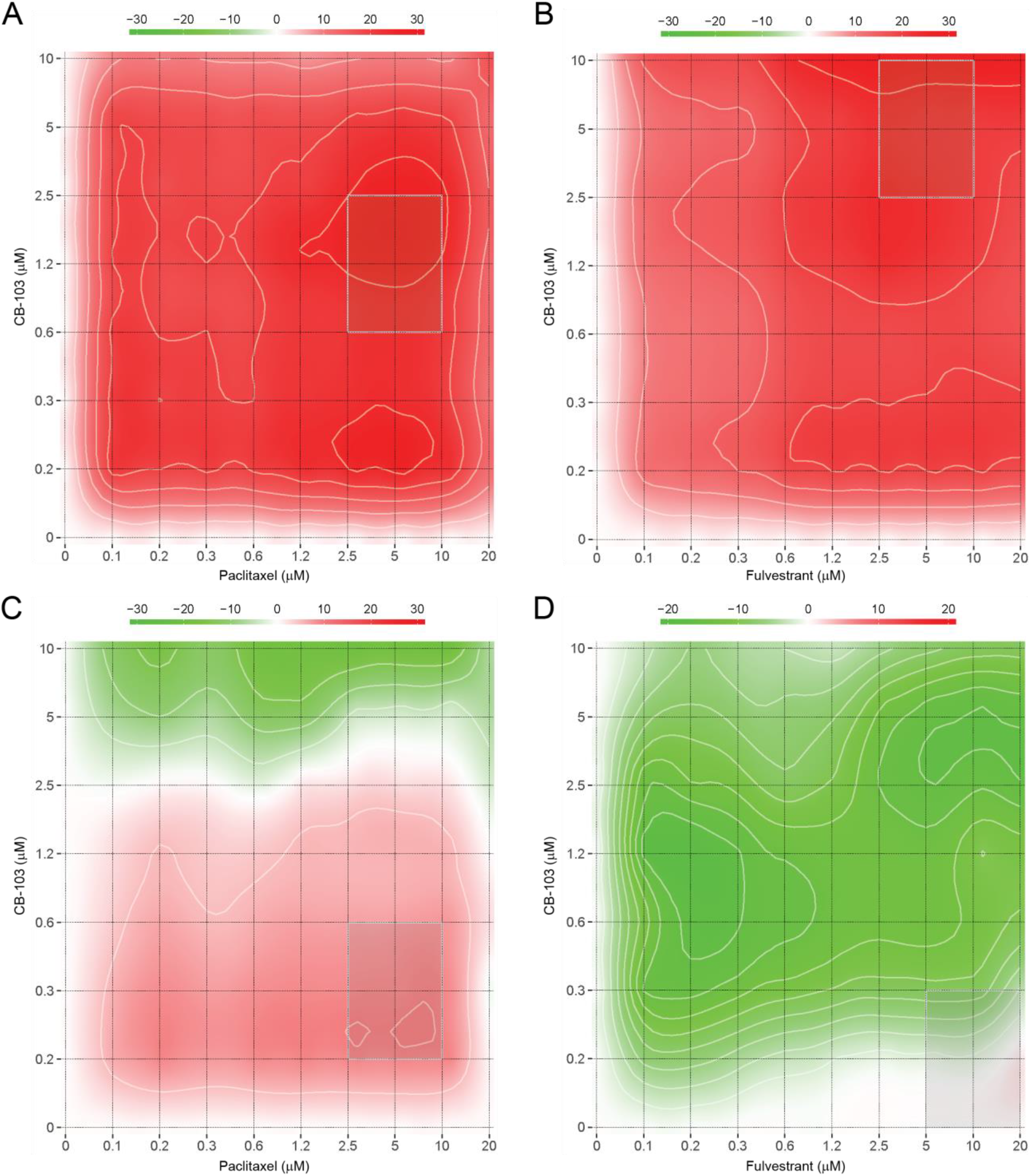
– CB-103 shows synergy with paclitaxel and fulvestrant in breast cancer cell lines. TNBC (HCC1187) and ER+ BC (MCF-7) cells were treated with combinations of CB-103 (150 nM – 10 mM) with paclitaxel or fulvestrant (75 nM – 20 mM) and cell viability was measured after 72 hours of treatment. Synergy between CB-103 and paclitaxel or fulvestrant in MCF-7 (A, B) and in HCC1187 (C, D) cells is reported as heatmap. >0, synergistic effect; 0, no combination effect; <0, antagonist effect.

### CB-103 strongly inhibits mammosphere formation when combined with fulvestrant

CSC in ERα+ and TNBC have been reported to be Notch-dependent (6, 16, 27, 28). Mammosphere formation assays are a useful surrogate for interrogating CSC activity in tumor cells. We sought to determine whether CB-103 inhibits mammosphere forming ability in an estrogen-dependent, endocrine-resistant cell line models in the presence or absence of fulvestrant. The choice of fulvestrant was supported by the combination screens described above and by mechanistic considerations. We previously showed that Notch1 can induce ERα-dependent transcription in the absence of estrogen (19). Notch1-induced transactivation of ERα target genes in the absence of estrogen still requires the presence of ERα, which is degraded in response to fulvestrant. Estrogen deprivation, mimicking aromatase inhibitors, leaves ERα intact and able to be activated by the Notch1-dependent mechanism we described (19). Notch4 NICD has the same activity (Singleton at al., in preparation). Mammospheres generated from estrogen-sensitive MCF-7 and two other endocrine resistant cell lines were treated with fulvestrant, CB-103 alone or combinations thereof. CB-103 at clinically achievable concentrations suppressed mammosphere formation alone and in combination with fulvestrant in MCF-7 (Figure 2A) and in the ER+ endocrine-resistant models T47D/PKCα (22) (Figure 2B) and WHM20, a PDX-derived cells expressing mutant ESR1-Y537S (23) (Figure 2C). Both these endocrine-resistant models have been reported to be Notch4-driven (19). In all cases, the combination of fulvestrant and CB-103 caused superior inhibitory effect on mammosphere formation compared to single agents. In these models, 1 and 5 μM CB-103 appeared to be equipotent. CDK4/6 inhibitors, such as Palbociclib (Pfizer, USA) are used as 2^nd^ line agents in endocrine-resistant ER+ tumors in combination with fulvestrant (29, 30). Palbociclib showed synergy with CB-103 in MCF-7 cells (Table 1). In WHM20 cells, carrying Y537S mutated ESR1, the combination of CB103 and palbociclib was significantly more potent in reducing mammospheres formation (Figure 2D) than CB-103+ fulvestrant (Figure 2C).

**Figure 2.**
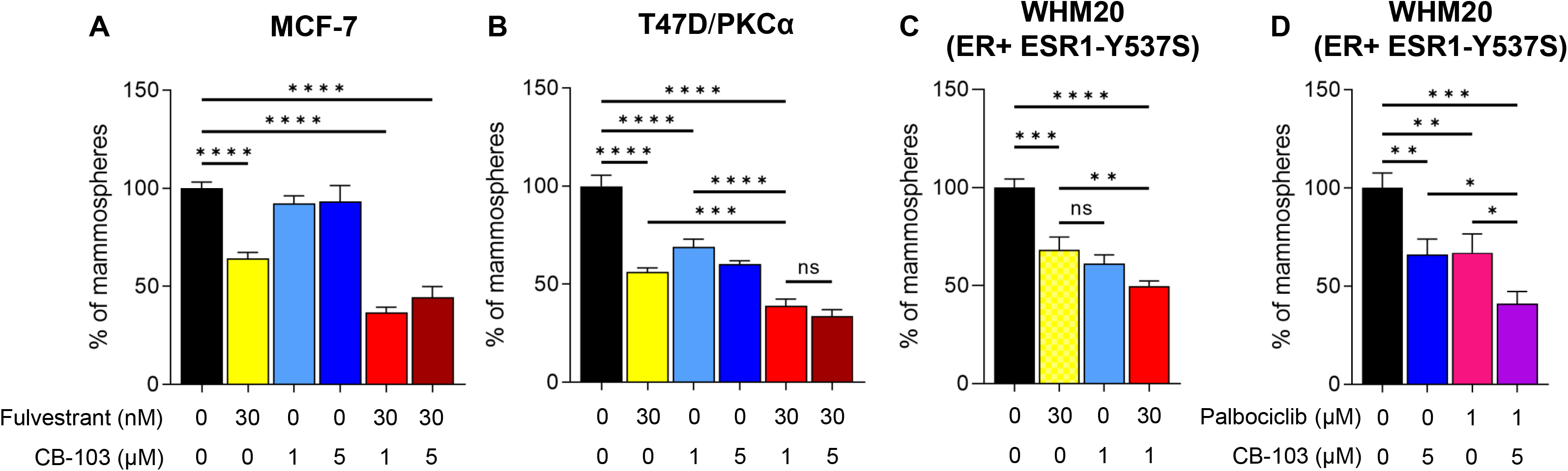
– CSC suppression by CB-103 alone or in combination second line endocrine therapies. CB-103 suppresses mammosphere formation alone and in combination with fulvestrant in wild type MCF-7 cells (A) and two distinct endocrine-resistant models, T47D/PKCα (B) and WHM20, PDX-derived cells expressing mutant ESR1-Y537S (C). In the ESR1 Y537S mutant line (D) CB-103 plus Palbociclib also shows potent inhibition of mammosphere formation. The concentration of each drug used is indicated under each bar according to x-axis legend. Statistical significance upon one-way ANOVA with Bonferroni correction for multiple comparisons is shown on each histogram with *=p<0.05; ** = p<0.01; *** = p<0.001; **** = p<0.0001.

### Combination therapy with CB-103 plus fulvestrant induces tumor regression in PDX derived, ESR1-mutant WHM20 cell line

CB-103 formulation was optimized for preclinical efficacy studies in mice in order to prolong exposure time and have a stable plasma concentration while lowering the highest plasma concentration (C_max_) responsible for the maximum tolerated dose (MTD). Indeed, orally or intraperitoneally administered CB-103, used previously in Lehal et al. 2020, has a shorter half-life in rodents with a very high C_max_. This fast release formulation appeared to have contributed to MTD in rodents. Therefore, we developed a subcutaneous formulation that allows slow release of the drug in the bloodstream. Pharmacokinetics (PK) analysis of a single dose of CB-103 60 mg/kg administered subcutaneously (s.c.), showed that the compound reached its maximal plasma concentration (C_max_) 1 hour post dose with sustained exposure up to 12 hours post dose (Additional Figure 1A). Moreover, plasma concentration upon single dose was comparable with CB-103 concentrations used during *in vitro* studies.

We performed pilot dose-finding experiment in mice by interrogating the reversible suppression of Marginal Zone B (MZB) cells as a validated pharmacodynamics (PD) biomarker for on-target Notch inhibition in animal model (20). We used non-tumor bearing nude mice to evaluate MZB inhibition. As shown in Additional Figures 1B and 1C, CB-103 led to a dose-dependent decrease in MZB cell population without apparent toxicity (Additional Figure 1D). This further validated the on-target activity of both doses of CB-103 *in vivo* in our model.

In a first efficacy study, we investigated a patient-derived ER^+^ BC xenograft harboring wild type ESR1 (CTG-2308). Since CB-103 60 mg/kg is the maximum tolerated dose, we combined a suboptimal dose of CB-103 (40 mg/kg) with fulvestrant. The combination of CB-103 and fulvestrant showed significant tumor growth delay compared to control group but the effect was comparable to the monotherapies (Figure 3A), consistent with the fact that wild type ESR1 is fully sensitive to fulvestrant.

**Figure 3.**
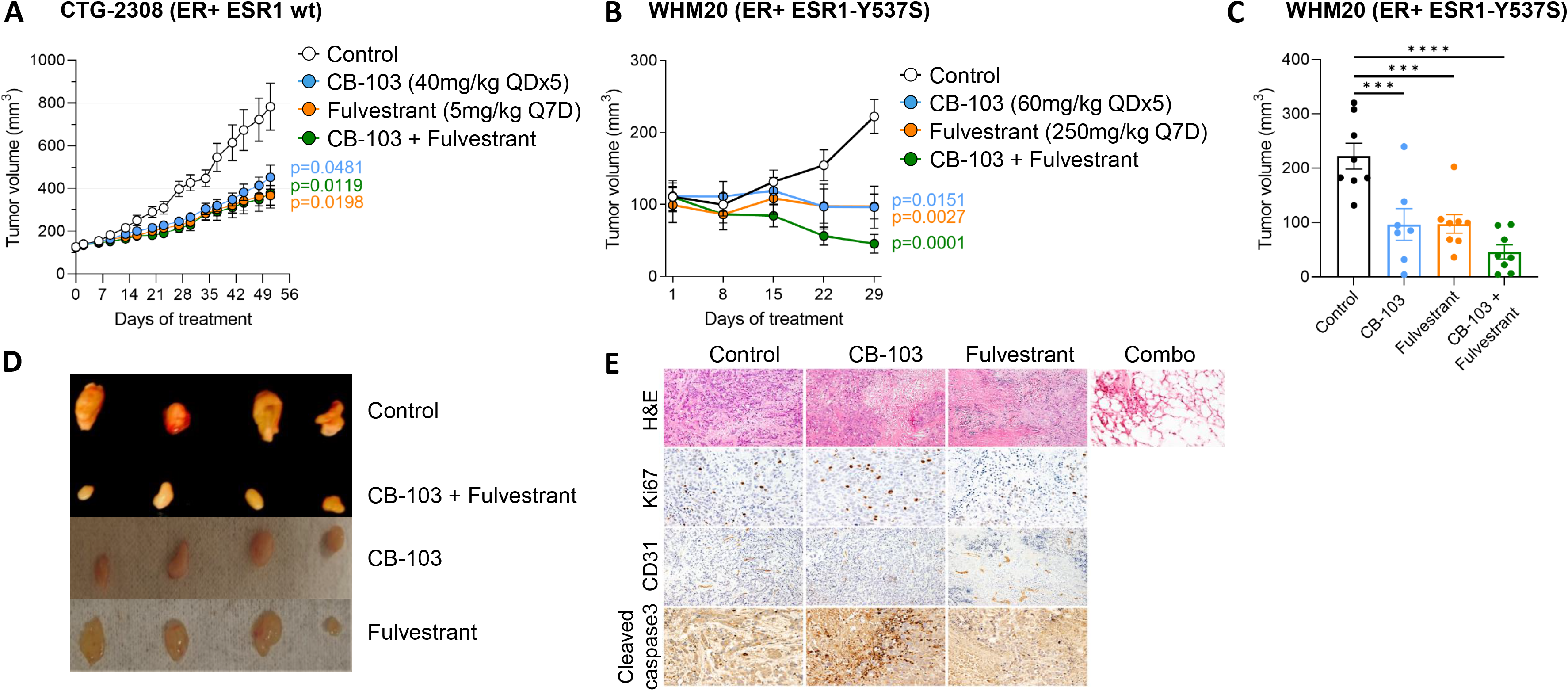
– Combination of CB-103 and fulvestrant inhibits the growth of an ESR1-mutant breast cancer model. A) Athymic nude female mice (n=10/group) engrafted subcutaneously with PDX CTG-2308 model (ER+ ESR1 wild type) were treated for 7 weeks with vehicle control, CB-103 (40 mg/kg QDx5) or fulvestrant (5 mg/kg Q7D) or their combination as shown. B) Athymic nude female ovariectomized mice (n=8/group) engrafted orthotopically with PDX WHM20 model (ER+ ESR1-Y537S) were treated for 4 weeks with vehicle control, CB-103 (60 mg/kg QDx5) or fulvestrant (250 mg/kg Q7D) or their combination as shown. C) Histogram plot of tumor volumes in Fig 3B on day 29 at the end of the treatment. Each bar represents Mean ± SEM of individual values identified by dots. *** = p<0.001; **** = p<0.0001. D) Picture of representative excised tumor samples (n=4) at endpoint from WHM20 cohort. E) Immunohistochemistry analysis of excised tumors. From top to bottom: H&E staining, proliferation marker (Ki67), anti-angiogenesis marker (CD31) and apoptosis marker (cleaved caspase-3). Only H&E staining was performed in tumor samples treated with combination.

We then explored the efficacy of CB-103 in an endocrine-resistant model. We selected PDX-derived WHM20 cells expressing mutant ESR1-Y537S (23) because of this model’s partial sensitivity to fulvestrant (23). We compared CB-103 side-by-side with the standard of care fulvestrant, as monotherapy or in combination in nude/ovariectomized mice. We used ovariectomized mice and no estrogen pellet for tumor engraftment with Y537S cell line to recapitulate the hormonal environment of post-menopausal women, who most often develop endocrine therapy resistant, recurrent or metastatic HR+ BC. Figure 3B shows that single agent treatment with either CB-103 or fulvestrant produced a “stable disease” outcome. Combination treatment caused significant tumor regression (Figures 3B, 3C and 3D), which in 2/8 mice led to “complete response” (no measurable disease). Blinded histological examination of excised tissue in mice showing “complete responses” was performed by a board-certified pathologist. Samples harvested from combination-treated mice showed no evidence of disease, with inflammation, necrotic tissue and fibrofatty tissue (Figure 3E, top row). CB-103 monotherapy caused apoptosis (as determined by cleaved caspase 3 staining) and anti-angiogenesis (as determined by CD31 staining, Figure 3E) consistent with described effects of Notch inhibition in breast cancer models (31, 32) and breast cancer patients (33, 34). Ki67 appeared modestly increased in tumors treated with CB-103 alone, which is consistent with the mechanism of apoptosis being via mitotic catastrophe, as previously showed (32). No immunohistochemistry other than H&E staining was performed in tumor samples pertaining to combination treatment (Figure 3E, top row) since residual tumor tissue was absent or barely sufficient for RNA extraction (see below) with combination treatment. No overt toxicity, weight loss or diarrhea were observed in treatment arms (weight chart shown in Additional Figure 1E).

### RNAseq and differential gene expression

We performed whole transcriptome RNA sequencing on excised tumor tissues from combination treatment, single agent treatments and vehicle control-treated tissues. The heat map generated from combination treatment vs untreated control tumors (Figure 4A) showed significant transcriptome differences among the treatment arms. Figure 4B depicts the Venn diagram of transcripts significantly affected by different treatment regimens. Four genes were common to all treatment groups, 11 genes were unique to CB-103-containing treatments only and 80 genes were modulated exclusively by the combination treatment. Importantly, when pathway analysis was performed, the estrogen signaling pathway was significantly impacted only in tumors treated with the combination (Table 2) and not in tumors treated with either single agent, consistent with our working hypothesis that combined inhibition of ERα and Notch is required to block ERα-dependent transcription in Notch-expressing endocrine-resistant tumors.

**Figure 4.**
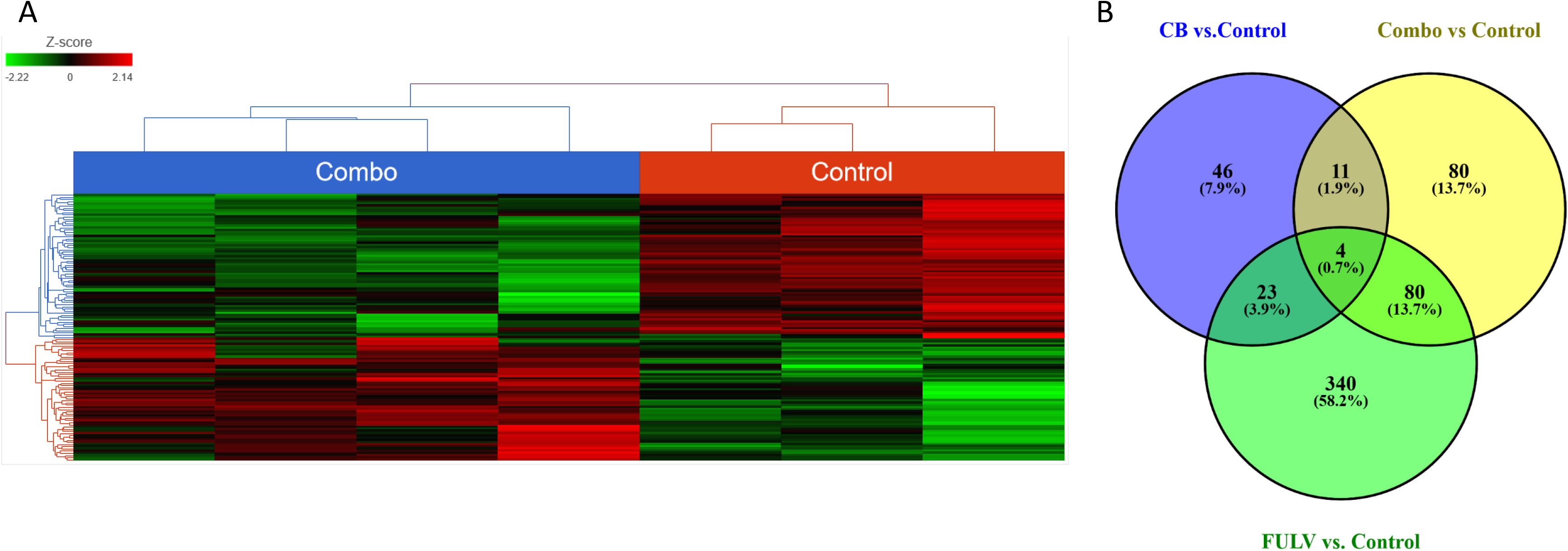
– Combination and control tumors present different gene expression patterns. A) Heat map for untreated control vs combination treatment was generated using FDR < 0.05 and fold change >2. B) Venn diagram depicting the number of common and unique genes in the various treatment arms; monotherapy and combination therapies (compared to the untreated control). Analysis adjusted for multiple comparisons using FDR < 0.05 and fold change >2.

**Table 2.** – Top 5 pathways modulated by each treatment vs control (IKGG).

| Relevant pathway enrichment | Enrichment score | p value |
| --- | --- | --- |
| <u>Control vs Fulvestrant</u> |  |  |
| cAMP signaling pathway | 10.6428 | 2.39E-05 |
| TGF-β signaling pathway | 8.06829 | 0.000313 |
| Complement and coagulation cascades | 6.78715 | 0.001128 |
| Calcium signaling pathway | 6.60461 | 0.001354 |
| cGMP-PKG signaling pathway | 6.35896 | 0.001731 |
| Aldosterone synthesis and secretion | 5.47526 | 0.004189 |
| <u>Control vs CB-103</u> |  |  |
| TGF-β signaling pathway | 4.36199 | 0.012753 |
| Bladder cancer | 3.91465 | 0.019948 |
| Arginine and proline metabolism | 3.69323 | 0.024891 |
| Drug metabolism – other enzymes | 3.22466 | 0.039770 |
| Transcriptional misregulation in cancer | 3.02111 | 0.048747 |
| <u>Control vs Combo</u> |  |  |
| Estrogen signaling pathway | 8.35942 | 0.000234 |
| Complement and coagulation cascades | 7.10170 | 0.000824 |
| Platelet activation | 5.41271 | 0.004460 |
| Cytokine-cytokine receptor interaction | 4.75188 | 0.008635 |
| cAMP signaling pathway | 4.51626 | 0.01093 |

### CB-103 induces a durable response in combination with Paclitaxel in TNBC

As stated earlier, HCC1187 is resistent to γ-secretase inhibition due to a Notch2 chromosomal translocation leading to constitutive activation of Notch signaling (20). Newman et al. (35) described all the mutations and the timing of their onset in TNBC HCC1187. Tumors produced by this Notch2-mutated cell line are resistant to GSIs and to Notch-blocking monoclonal antibodies (25). We have previuously shown that CB-103 treatment alone potentily inhibited tumor growth in HCC1187 tumors in nude mice compared with control (25). Lehal et al. (25) further validated the specificity of target engagement by CB-103 as demonstrated by the downregulation of Notch target gene HES1, whereas GSIs such as DAPT and RO4929097 didn’t induce any noticeable change in HES1 in this model. Given that GSIs have shown acceptable safety in breast cancer patients combinations including paclitaxel (36) and CB-103 has a better toxicity profile than GSIs, we explored whether the synergy we observed *in vitro* between paclitaxel and CB-103 translates into *in vivo* efficacy without undue toxicity. We interrogated whether combining CB-103 with standard of care chemotherapeutic agent paclitaxel would modulate tumor growth differently. Figure 5A demonstrates that continuous single-agent treatment with CB-103 (60 mg/kg QDx5) for two weeks induced 32% tumor growth inhibition (TGI) compared to vehicle treated arm. Paclitaxel (10 mg/kg Q7D) alone or in combination with CB-103, on the other hand, induced 102% and 104% TGI. At first glance, the incorporation of CB-103 in combination treatment arm didn’t seem to offer any additional benefit over paclitaxel monotherapy. When vehicle group reached endpoint, treatment in the other dosing groups were discontinued while mice were still monitored for about four weeks (Figure 5B). Strikingly, after 14 days post treatment discontinuation, tumor growth rebounded immediately in the paclitaxel treated arm, whereas in the combination treatment arm tumor growth was delayed by 12 days. The survival graph in Figure 5C illustrates the benefit of combination treatment vs paclitaxel alone.

**Figure 5.**
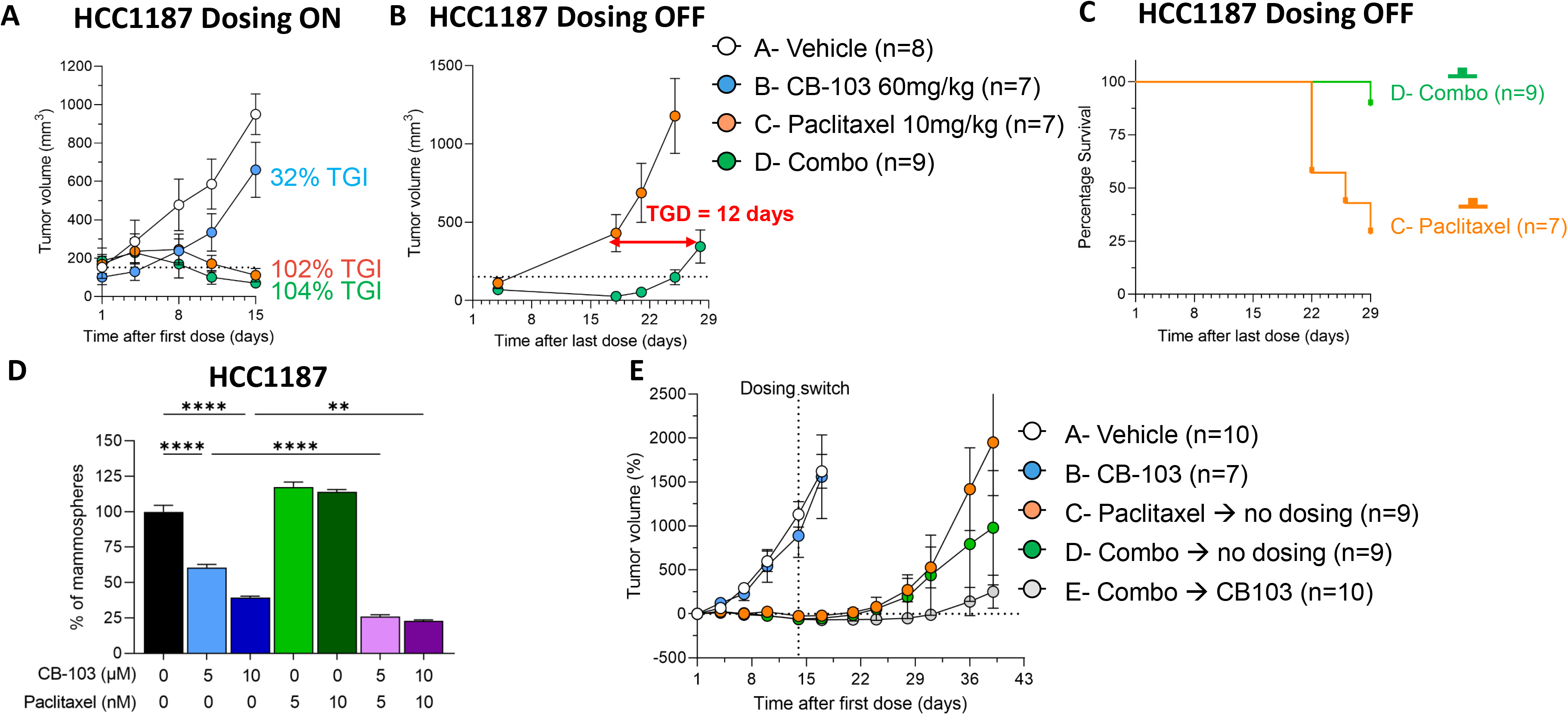
– CB-103 causes a durable response in combination with Paclitaxel in a TNBC model. A) NSG immune compromised female mice engrafted subcutaneously with TNBC cell (HCC1187) were treated for 2 weeks, before control group reached endpoint, with CB-103 (60 mg/kg QDx5) or paclitaxel (10 mg/kg Q7D) or their combination. At control group endpoint all dosing was discontinued, and tumor relapse (B) and animal survival (C) were still monitored for around 4 weeks in order to determine tumor growth delay (TGD). D) Mammosphere culture was established from HCC1187 cells and treated for 7 days with the indicated concentration of CB-103 or paclitaxel. Each bar represents Mean ± SEM of individual values identified by dots. ** = p<0.01; **** = p<0.0001. E) Mice engrafted subcutaneously with TNBC cell HCC1187 were treated for 2 weeks, before control group A reached endpoint, with CB-103 (60 mg/kg QDx5) or paclitaxel (10 mg/kg Q7D) or their combination. On dosing day 14, treatment in groups C, D and E was switched as indicated in figure legend and tumor relapse was monitored until group C reached endpoint.

Next, we examined whether the delayed tumor growth in the combination-treated arm may have been due to CSC inhibition. Mammosphere formation assays in HCC1187 cells treated with CB-103, paclitaxel and the combination thereof (Figure 5D) were performed. In agreement with our observation in endocrine-resistant ER+ cell lines, we observed significant decrese in the mammosphere forming ability of HCC1187 cells treated with CB-103. Combination treatment with 5 nM paclitaxel plus 5 μM CB-103 induced potent suppression. Increasing the dose of CB-103 and Paclitaxel to 10 μM and 10 nM respectively didn’t further increase this effect. Notably, treatment with paclitaxel alone didn’t reduce mammosphere forming ability. This is consistent with previous findings by Tanei et al. (37), who didn’t notice any efficacy of Ixabepilone, a microtubule-targeting chemotherapeutic drug, on mammosphere forming ability in the MDA-MB-231 TNBC cell lineIn an independent experiment we sought to examine whether continuous CB-103 treatment of TNBC tumors following paclitaxel treatment discontinuation significantly delays tumor growth in mice (Figure 5E). Figure 5E, arm E shows that continuous dosing with CB-103 alone for up to 25 days after combination treatment for two weeks significantly delayed tumor growth in mice, whereas, tumor growth rebounded relatively quickly following a two week combination traetment (Figure 5E, arm D). Interestingly, we determined that the efficacy of combination of CB-103 and paclitaxel is similar when applied to small or large volume tumors (Additional Figure 2). Indeed, when the combination therapy was applied to mice with average tumor volume of 50 mm^3^ or 500 mm^3^, the slope of tumor regression was the same (Additional Figure 2B). Since paclitaxel exposure predisposes subjects to more toxicities compared with CB-103 monotherapy, treatment with CB-103 as a maintenance therapy following combination treatment may offer new opportunities in the clinical setting. CB-103 has a favorable safety profile compared to paclitaxel and doesn’t induce GI toxicities associated with GSIs. (18, 20, 25).

## Discussion

Our data provide preclinical proof of concept for the investigation of CB-103, a first-in-class transcriptional pan-Notch small molecule inhibitor with superior safety profile compared to GSIs (6, 18) in two clinically challenging breast cancer subtypes, namely, endocrine resistant HR+ cancers and TNBCs.

A significant fraction of patients with ER+ BC treated with endocrine agents will see their disease relapse. Multiple molecular mechanisms have been described as drivers of endocrine resistance. Development of *ESR1* mutations, activation of several signaling pathways such as PI3K or MAPK or loss of ER expression are among the most commonly observed resistance mechanisms (6). This heterogeneity in mechanisms of resistance can be also drug-specific, as shown by *ESR1* mutations that confer resistance to aromatase inhibitors but not to selective estrogen receptor degraders (SERDs) (38). Several groups (39–41) independently showed that mutations in the ligand binding region of the estrogen receptor alpha gene (ESR1) can cause endocrine resistance. The development of *ESR1* mutations (6, 41) is one of the most common acquired endocrine resistance mechanisms in ER+ BC. Fulvestrant remains a standard of care for post-menopausal HR+ women who fail to respond to AI plus CDK4/6 inhibitors. However, for most patients who respond to fulvestrant, tumors eventually recur, and third line therapies are largely ineffective. CB-103 showed strong synergism with fulvestrant and with palbociclib in an unbiased pharmacological screen and suppressed mammosphere formation alone and in combinations with either fulvestrant or palbociclib in different endocrine therapy resistant ER+ cell lines. In vivo, CB-103/fulvestrant combination treatment caused tumor regression and was superior to either agent alone in an ESR1-Y537S mutant model, while CB103 monotherapy or the doublet combination had comparable efficacy to fulvestrant alone in an ESR1 wild-type model. KEGG pathway analysis of tumor RNASeq data showed that the ER signaling pathway was the most impacted pathway in ESR1-Y537S mutant tumors treated with the combination but not with either single agent (Table 2). This is consistent with our hypothesis that combined inhibition of ERα and Notch is required to block ERα-dependent transcription in Notch-expressing endocrine-resistant tumors. Notably, in ER^+^ BC, ligand-independent Notch activation is also induced in the CSC population by sphingosine 1-phosphate through sphingosine 1-phosphate receptor 3 (S1PR3), and sphingosine kinase 1-positive CSC were highly tumorigenic (6). The fact that CB-103 can block both ligand dependent and ligand-independent Notch activation offers a clear benefit compared to Notch inhibitors targeting ligands. We propose that future clinical trials of combinations between Notch inhibitors and endocrine therapy should explore selective SERDs rather than AI, at least in part because Notch1 can induce ERα-dependent transcription in the absence of estrogen (11). Our results suggest that ESR1-Y537S mutation may be a biomarker for sensitivity to CB-103/SERD combinations in humans, consistent with the observations of Gelsomino et al. (13). Mammosphere results from this study and our previously published data (19) (see below) suggest that PKCα expression may also be a biomarker for sensitivity to this combination. We saw no indication of intestinal toxicity in CB-103 treated mice, as monotherapy or with fulvestrant. We had previously shown that when combined with estrogen deprivation or endocrine therapy, the intestinal toxicity of intermittently administered GSIs in preclinical models was greatly ameliorated without compromising efficacy (42). These findings were confirmed in the clinical setting. In a pilot phase 1b clinical trial, we showed that since discontinued GSI RO4929097 in combination with exemestane in metastatic or relapsed, endocrine-resistant breast cancer was well tolerated for up to 6 months, without significant intestinal toxicity. Clinical responses were observed in 8/14 evaluable patients. These include one partial response (PR), 7 stable diseases (SD) and 6 progressions of disease, but not complete responses (CR). The total clinical benefit rate (CR+PR+SD ≥ 6 months) was 20%, with a progression-free survival (PFS) of 3.2 months. In the same study, (19) we showed that mammospheres generated from T47D:PKCα cells were partially sensitive to fulvestrant as a single agent (19). GSI RO4929097 alone (5 μM) significantly decreased mammosphere numbers, consistent with the results of Simoes et al. (43). The combination of GSI plus fulvestrant was significantly more potent than either agent alone, nearly abolishing mammosphere growth (19). Our results indicate that CB-103 has a similar activity profile to GSIs, with no apparent toxicity at the doses we evaluated, and suggest that at least part of the anti-neoplastic effect of CB-103 in breast cancer models is due to anti-CSC activity (19). This is consistent with the literature: treatment with tamoxifen or fulvestrant in patient-derived samples and xenograft models of ER+ breast tumor selected CSC-like cells through upregulation of the Jagged1-Notch4 signaling axis (18). Combination treatment with Notch inhibitors reduced the frequency of hormonal therapy-resistant CSC (18).

Selective CDK4/6 inhibitors (CDKis) (44) are FDA-approved in combination with endocrine therapy to treat patients in first and second-line settings (29, 30, 45). CDKis induce G1 cell cycle arrest by preventing the phosphorylation of the retinoblastoma (Rb) protein. There is clear mechanistic rationale for exploring combinations of Notch inhibitors with CDKis. Notch1 and Notch4 can drive cell cycle progression in the absence of estrogen in ERα-positive breast cancer cells and induce expression of cyclins A and B (10). Mitotic kinases CDK1 and 2 phosphorylate the active intracellular form of Notch1, leading to its degradation (46). This suggests that cell cycle arrest at the G1S stage caused by CDK inhibitors, which prevents mitosis, would lead to accumulation of cleaved Notch1 and possibly CDK inhibitor resistance through upregulation of cyclins D1 (11), A and B (10). Our data indicate superior inhibitory efficacy of palbociclib/CB-103 combination compared with palbociclib/fulvestrant in mammosphere formation assays in ESR1 mutant Y537S cells. This suggests that CB-103 could be clinically explored in doublet combinations with CDKis without SERDs. We did not explore triple combinations in this study, as a CB-103/fulvestrant combination was sufficient to cause complete responses in vivo. Future experiments will explore the effectiveness of triple combinations and CDKi/CB-103 doublets. Taken together, our data support the notion that Notch inhibition in combination with standard of care has potential clinical interest in endocrine-resistant ER+ tumors carrying the ESR1 Y537S mutation and possibly other biomarkers associated with increased Notch activity (e.g., PKCα expression).

We and others have shown that Notch signaling plays a significant role in TNBC. Notch1 expression portends to poor survival in recurrent TNBC (47). There is extensive literature supporting the notion that active Notch signaling and expression of Notch ligand Jagged1 are biomarkers of poor prognosis and potential therapeutic targets in TNBC (48–54). Of note, Notch signaling cooperates with EZH2, a druggable target, in TNBC (55) and represses PTEN in this group of tumors (52). Furthermore, Notch signaling promotes treatment-resistant CSC in TNBCs treated with TORC1/2 inhibitor (16). Notch1 inhibition increases sensitivity to paclitaxel in a breast cancer model by affecting the pool of CSC through miR-34a upregulation (56). TNBC cell lines with *Notch1*-activating mutations were shown to be sensitive to GSI MRK-003 alone and in combination with paclitaxel (18). TNBC cells with *Notch2* rearrangement are GSI-resistant (20). Our results provide a rationale for considering CB-103 as an attractive option for a clinical trial in TNBC, in combination with a taxane-based standard of care chemotherapy regimen. Initial trials may focus on patients with *Notch1* or *Notch2* rearrangements.

## Conclusion

Our data indicate that CB-103, a second generation, clinical stage transcriptional Notch inhibitor, is an attractive candidate for clinical investigation in endocrine resistant, recurrent breast cancers with biomarker-confirmed Notch expression in combination with SERDs and/or CDKis and in TNBCs with biomarker-confirmed Notch expression in combination with taxane-containing chemotherapy regimens.

## Declarations

### Ethics approval and consent to participate

Approval was obtained for all animal studies under the guidelines of IACUC as mandated by LSUHSC and under Swiss National license n. 33520, Cantonal license n. VD3672.

### Consent for publication

Not applicable

### Availability of data and materials

All data generated or analyzed during this study are available from the corresponding author on reasonable request.

## Authors’ contributions

LM and RL conceived and designed the study; SM, MV, CU, SL collected and analyzed data; SM, LM, RL drafted the manuscript; SM, MV, CU, LM, RL interpreted data and substantially revised the manuscript. RR helped design experiments, JZ, GM, FH, LD generated data; JZ, LD generated figures.

## List of Abbreviations

ANOVA: Analysis of variance
BC: Breast Cancer
CDK: Cyclin-dependent kinase
C_max_: Maximal plasma concentration
CR: Complete response
CSCs: Cancer Stem Cells
DMSO: Dimethyl sulfoxide
ER+: Estrogen receptor positive
ESR1: Estrogen Receptor Gene α
FCS: Fetal Calf Serum
GSIs: gamma-secretase inhibitors
H&E: Hematoxylin and Eosin
LC-MS: Liquid chromatography–Mass Spectrometry
MAPK: Mitogen-activated protein kinase
MZB cells: Marginal Zone B cells
PD: Pharmacodynamics
PDX: Patient-derived xenograft
PI3K: Phosphoinositide 3-kinase
PK: Pharmacokinetics
PR: Partial response
PR+: Progesterone receptor positive
SD: Stable disease
SERD: Selective Estrogen Receptor Disruptor
TNBC: Triple Negative Breast Cancer
TV: Tumor volume

## Acknowledgements

We sincerely acknowledge the technical supports from Translational Genomic core (TGC), Cellular Immunology and Immune Metabolism Core (CIMC) and Molecular Histopathology and Analytical Microscopy Core (MHAM) of LSUHSC.

## Figure titles and legends

**Additional Figure 1.** – A) Plasma concentration of CB-103 at 0.25, 0.5, 1, 2, 4, 8, 12 hours after single dose at 60 mg/kg. Each dot represents mean ± SEM of n=3 mice. B) Dose dependent downregulation of Marginal Zone B (MZB) cells in mice upon 7 days of CB-103 daily dosing; flow cytometry data showing percentage of MZB cells of representative mouse from vehicle control, 40 mg/kg and 60mg/kg of CB-103 treatment; C) Histogram plot of the mean ± SEM frequency of MZB cells for each dosing group (n=4). D) Weight plot of mice in the target engagement cohort in the MZB cell experiment. E) Weight chart of mice in the Fulvestrant efficacy cohort indicating tolerability of doses. F) Weigh chart of mice in the paclitaxel efficacy cohort indicating tolerability of doses.

**Additional Figure 2.** – Efficacy of CB103 + paclitaxel therapy on small and large TNBC tumors. NSG immune compromised female mice engrafted subcutaneously with TNBC cell line HCC1187 were treated with combination therapy (CB-103 60 mg/kg QDx5 + Paclitaxel 10 mg/kg Q7D) once average tumor volume reached 50 mm^3^ (green dots) or 500 mm^3^ (white dots). Tumor growth was evaluated by measuring tumor volume (A) or the percentage difference of volume from beginning of dosing (B).

