## Supplementary figures and images for "The efficacy of CB-103, a first-in-class transcriptional Notch inhibitor, in preclinical models of breast cancer"

### Supplemental Figures

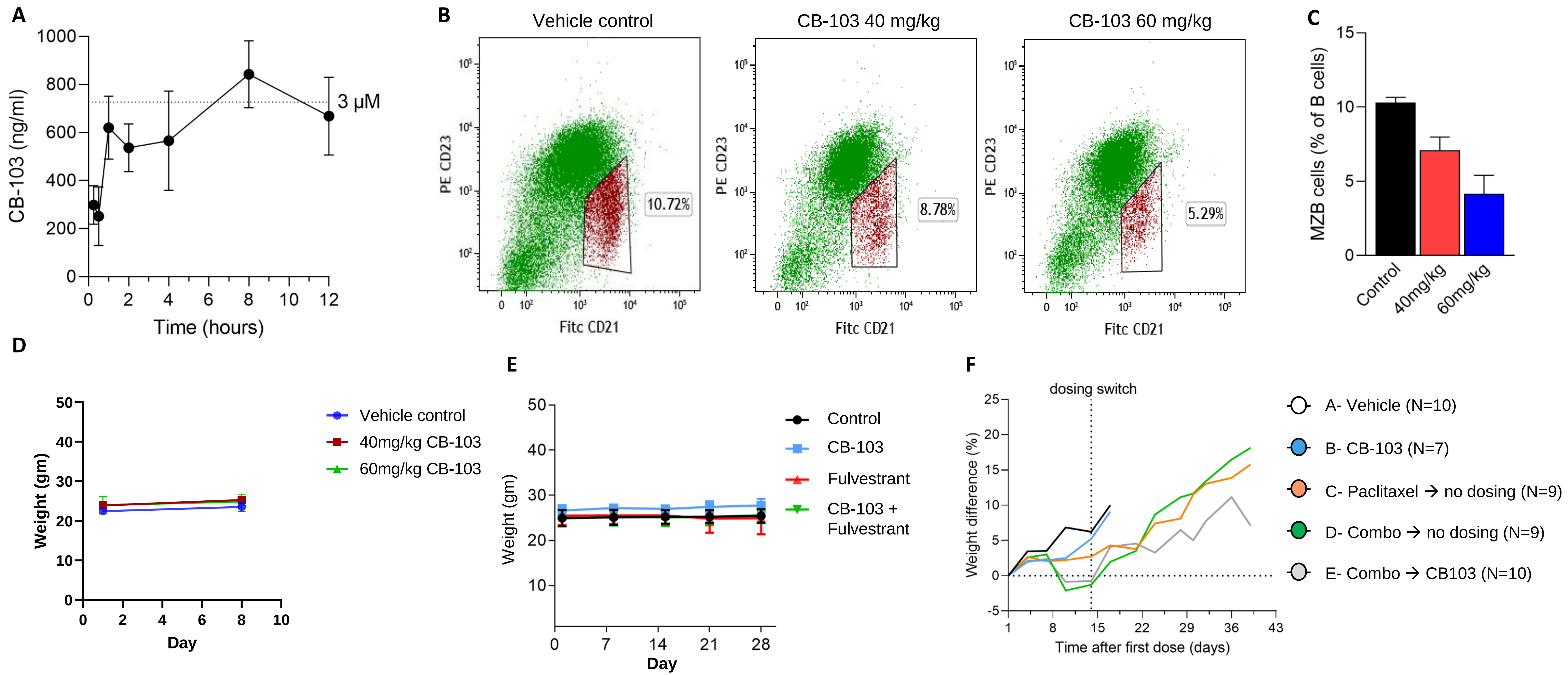

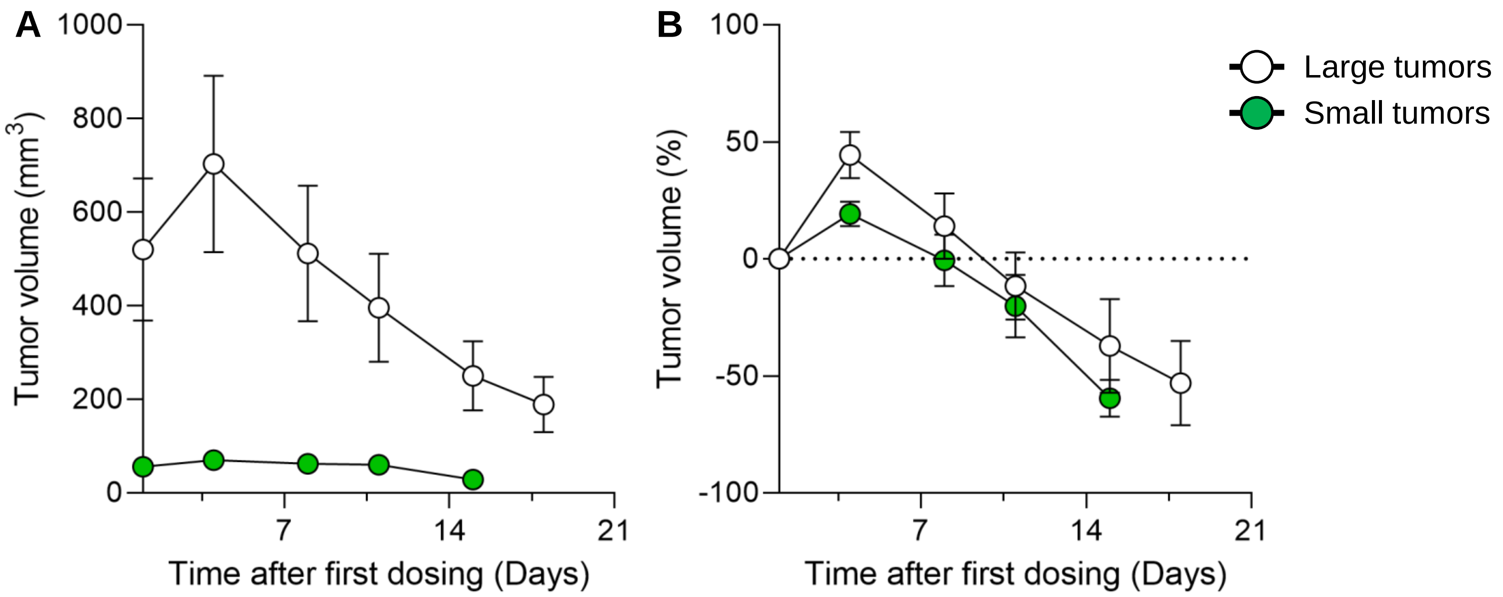
